# High-Resolution Time-Lapse Imaging of Droplet-Cell Dynamics via Optimal Transport and Contrastive Learning

**DOI:** 10.1101/2025.03.06.641128

**Authors:** Luca Johannes Schlotheuber, Michael Vollenweider, Sven Gutjahr, Tiago Hungerland, Richard Danis, Weronika Ormaniec, Aline Linder, Valentina Boeva, Ines Lüchtefeld, Klaus Eyer

**Affiliations:** Laboratory for Functional Immune Repertoire Analysis, Institute of Pharmaceutical Sciences, Department of Chemistry and Applied Biosciences, ETH Zürich, Zürich, Switzerland; Department of Computer Science, Institute for Machine Learning, ETH Zurich, Zurich, Switzerland; ETH AI Center, ETH Zurich, Zurich, Switzerland; Department of Biomedicine, Aarhus University, The Skou Building, Høegh-Guldbergs Gade 10, 8000 Aarhus C, Denmark

**Keywords:** Single-cell analysis, Droplet Microfluidics, Tracking, Optimal-Transport, Deep Contrastive Learning

## Abstract

Single-cell analysis is essential for uncovering heterogeneous biological functions that arise from intricate cellular interaction. Microfluidic droplet arrays enable precise dynamic data collection through cell encapsulation in picoliter volumes. The time-lapse imaging of these arrays can reveal functional kinetics and cellular fates, but accurate tracking of cell identities across time frames remains challenging when droplets move significantly. Specifically, existing machine learning methods often depend on labeled data or require neighboring cells as reference; without them, these methods struggle to track identical objects across long distances with complex movements. To address these limitations, we developed a pipeline combining visual object detection, feature extraction via contrastive learning, and optimal transport-based object matching, which minimizes reliance on labeled training data. Our approach was validated across various experimental conditions and was able to track thousands of water-in-oil microfluidic droplets over large distances and long (*>* 30 min) time-separated frames. We achieved high precision in previously untraceable scenarios, tracking small, medium and large movements (corresponding to ~126, ~800 and ~10,000 *µ*m, respectively) with a success rate of correctly tracked droplets of *>* 90% for average movements within 2-12 object diameters, and *>* 60% for average movements of *>* 100 object diameters. This workflow lays the foundation for high-resolution, dynamic analysis of droplets and cells in both spatial and temporal dimensions without relying on visual labeling, allowing high-accuracy tracking in samples, where the uniqueness of the sample makes repeating experiments infeasible.

## Introduction

In recent years, the development of single-cell technologies has grown immensely, allowing researchers to study cellular populations through the lens of heterogeneity and plasticity and discover new cell types and functions. While much work has been done on the transcriptomic level through sequencing approaches, RNA and DNA by itself are often a poor descriptor of protein functionality, that is often best assessed directly. Moreover, the time-component of such analysis is essential as cells display dynamic plasticity and changes over time. Therefore, imaging and tracking living cells is often a necessity.

Applications which employ such direct, dynamic analysis have been growing following breakthroughs in a wide range of research domains: from tracking cells inside living organisms [1], intra-vital imaging [2], [3], and monitoring of singlecells or organoids in lab-on-a-chip devices [4] to droplet based dynamic phenotyping of cell secretion inside droplets [5]. Central to all these methods is the ability to track single cells to correctly monitor their fates over time. While advanced tracking algorithms powered by machine learning have been developed, they often depend on visual cues and the relative positioning with respect to neighboring cells for accurate analysis [6]. Moreover, these algorithms are generally not optimized for “moving” systems and consistent spatial recovery [7]. Finally, available algorithms are often specialized for wells or tissues with small, fixed number of cells, rendering them unable to track in large images, thus lacking scalability and throughput.

To analyze protein secretion, droplet-based microfluidics has emerged as a transformative tool with many potential applications in modern medicine and fundamental science [8] [9] [10]. In essence, these technologies leverage the ability to isolate and analyze individual cells within microscopic droplets, efficiently linking the secreted molecules that are related to function of the cell that secreted them. Applying these technologies can offer new insights into cellular identity, functionality, behavior, and drug responses at a single-cell level. The power and potential of droplet microfluidics have been emphasized in recent studies, especially for the analysis of cell-cell interactions which highlight its application in high-throughput screening, diagnostics, and therapeutic development [8]. A subset of droplet microfluidics, 2D droplet microfluidics, has further emerged as an interesting proposition to study cellular functionalities, as this technique allows for measurement over time, allowing for the discrimination of sequential and simultaneous functionalities, cellular plasticity, cell-cell interactions, and fate [11] [12]. This advancement is crucial for understanding the dynamics of complex biological processes that occur on the level of individual cells, as it enables real-time observation of how individual cells interact and respond to different stimuli within a controlled microenvironment.

In practice, 2D single-cell analysis relies on the separation of cells into individual arrays, chambers of droplets, and the measurement over time [13] [14] [15] [16]. Here, uncontrolled movement over time, such as by larger droplets, air, imperfect filling of immobilization chambers, might result in the loss of correct data traces assigned to each cell. This is especially problematic when droplets move a distance larger than their radius, as the droplet centroid of the previous time-point is used as the main reference point for traditional methods, making them fail to track effectively. When there is movement greater than a certain percentage of the droplet radius between the different time points, data is often lost. In fact, experiments with movement *>* 10 pixel could not be tracked prior to our approach and off-the-shelve, commercial software applications for tracking this data are not available. The long intervals of acquisition (often needed for large and multi-spectral acquisition of images) further complicate this issue, allowing moving droplets to travel greater distances between frames. Finally, tracking through signal (inside cells or droplets) is problematic as it requires a biological understanding that might not be present during measurement and could be biased by the cellular heterogeneity and changing morphology of living cells. With the growing demand for larger-scale and deeper analytical resolution (*>* 10.000 events, *>* 10 parameters in parallel), time intervals and dataset sizes will continue to increase, underscoring the need for more advanced and comprehensive tracking approaches. Since droplet and cell images often display optical ambivalence, large and complex movement patterns without preserving local structures present a serious data analysis challenge.

Another challenge for 2D cell and droplet tracking experiments is that droplet microfluidics are created in a continuous flow from which the images are sampled. Lack of (physical) image boundaries means droplets can enter and exit between frames. In a not-fully immobilized system, this leads to variable droplet numbers across different frames. Recently developed or commercially available software struggle to resolve such data [5] [17]. Although cell line experiments can be repeated, potential data loss or inaccurate assignment due to movement presents a roadblock in the data analysis. This becomes crucial for rare, sensitive, or unique samples where it is essential to maintain high throughput and repetition may not be possible. With important clinical samples, inaccurate tracking can lead to incorrect assignment, producing false conclusions in the worst case. Only when cells are correctly tracked do these techniques enable the accurate extraction and resolution of relevant data that can be further used. Therefore, novel algorithms are needed to allow for the successful tracking of cellular functionality when movement is present in these arrays.

Given the potential uses of high-resolution dynamic singlecell data and the problems that droplet movements present for their analysis, this study introduces a novel approach for processing over-time single-droplet experiments. Our approach integrates positions and visual representations of the droplets into an optimal transport (OT) algorithm to enable effective tracking of the droplets across space and time. The pipeline we present consists of (i) a pre-processing, (ii) a droplet detection, (iii) a visual feature extraction, and (iv) finally a matching and uncertainty quantification step (Figure 1). We also provide a tool to visualize individual predicted trajectories with their respective certainty levels. We evaluate the visual feature extraction and show that our model produces meaningful features for the droplets, which stay consistent over time. Next, we evaluate the full pipeline heuristically on real data and quantitatively on simulated data. Finally, we link the algorithm to a read-out algorithm that calculates fluorescent mean intensities, which we use to track color-coded droplets across time and space and verify the predicted trajectories (Figure 1).

**Fig. 1:**
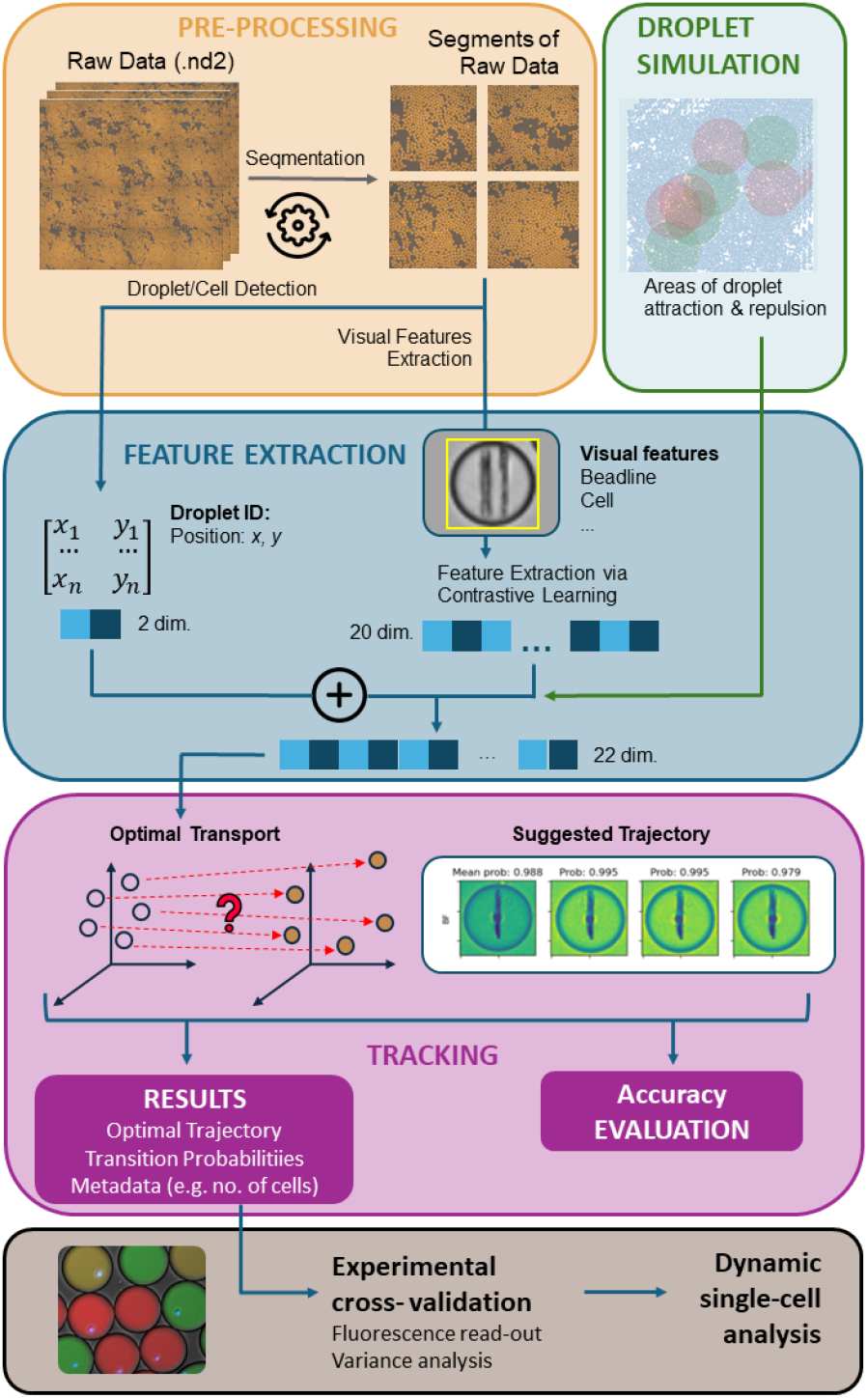
The main pipeline of data analysis from pre-processing to trajectory generation. Starting with raw-Data & segmentation; Simulation modeling for validating labeling and accuracy during (droplet population) movement. Different types of movement were simulated such as areas of attraction (droplets moving towards one position) and repulsion (droplets moving away); followed by Optimal transport (OT) implementation and trajectory generation, lastly from trajectories fluorescence read-outs are generated and validated enabling dynamic singlecell analysis.

## Methods

### (iv) Experimental time-lapse imaging data

Time-lapse epi-fluorescence imaging produces image tilesfield which are merged to a consecutive image spanning 10×10 mm typically. Generally, over-time experiments are acquired every 15, 30 or 60 minutes and contain between 5 and 15 intervals (Figure 3). In this study, samples were produced using a microfluidic chip as described by Linder et al. [12] [18]. Cells included peripheral blood mononuclear cells (PBMCs) donated from the Zürich blood bank and Chinese Hamster Ovarian (CHO) cells (Eurofins). Both experimental setups were produced in technical replicates with different degree of movement introduced. Specifically, **Dataset 1** (Linder et al. 2024) included experiments for the detection of cytokines from PBMCs after stimulation [18]. Bioassays are further described in [5] [12]. Such images include immune cells encapsulated with nanoparticles used for protein capture and fluorescent detection probes or fluorescent dye barcodes respectively. From this set, four experiments were run, differing in their total movement (based on Earth mover’s distance, EMD), ranging from small to large movement. The images of the droplet experiments were captured in up to 5 different channels including: DAPI, FITC, TRITC, Cy5 and a bright-field (BF) channel. The 2nd set of experiments, **Dataset 2**, consisted of images with droplets with CHO cells, without nanoparticles. Further, no adaption or fine-tuning of the contrastive learning or OT-problem balancedness was performed. Technical replicates with low-to-medium and large movements were performed from this data, referred to as biological crossvalidation dataset (Figure 7).

### B. Simulated data

Following our observations on experimental data, we simulated trajectories reflecting three different types of motion: divergence spots (droplets had a tendency to move away from red circles), attraction spots (droplets had a tendency to move into green parts), and big droplets that moved at large speed through the image as described in section III-B and IV-D. To have the droplets’ visual features, we used a data set of already tracked droplet trajectories and paired them with the simulated locations. Details of how the simulated data was integrated into the pipeline can be found in the Appendix IV; simulated data are available at https://github.com/ls154/DropletTracking/.

### (C) Pre-processing and object detection

Pre-processing involved image segmentation and subsequent droplet and cell detection. Large time-lapse image files (*>*10 GB) can strain computational resources; thus, we implemented an optional step to segment frames into disjoint segments, reducing memory requirements. We translated and implemented in Python the cell- and droplet detection algorithms from DropMap, a MatLab-based droplet detection and analysis software algorithm [5]. The implementation of the droplet-circle detection was adjusted to create an end-to-end pipeline importing all necessary parameters for Optimal Transport (OT)-based tracking. Specifically, we provided the option to detect droplets in the preprocessed bright-field channel using a combination of the Hough Circles algorithm [19] and the RANSAC (Random Sample Consensus) algorithm [20]. To avoid false-positives, droplets exiting image boundaries after measurement (in between time intervals) were not considered for tracking, while droplets entering were added continuously to the pipeline. A more detailed description can be found in the Appendix I and Supplemental Figure 1.

### (D). Visual feature extraction via contrastive learning

To incorporate visual features in the droplet tracking, we trained a convolutional neural network (CNN) to learn a lowdimensional vector representation of each droplet. As the CNN encoder backbone, we chose an EfficientNet-B1 model, due to its optimal features balanced between computational size of the model, accuracy, and efficiency [21]. The model was then fine-tuned to account for the tracking. The input to the encoder were droplet images of size 40 × 40 pixels with 2 channels, the preprocessed bright-field and DAPI cell channels. While the bright-field channel was used to obtain a visual representation of the beadline (crossing each droplet), the DAPI channel contained the stained cell. Finally, the output of the CNN embedding droplet images was a 20-dimensional feature vector:

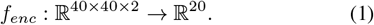

To produce meaningful representations, we used a contrastive learning approach to induce closeness in the latent space of two droplets from the same trajectory, while increasing distance in the latent space of two droplets from different trajectories. For this purpose we employed the InfoNCE loss function for a batch of representations Z as follows [22]:

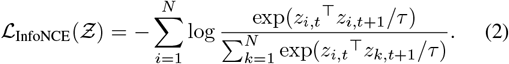

Here, *z*_*i,t*_ = *f*_*enc*_(*x*_*i,t*_) denotes the learned representation of droplet *i* at time step *t, N* is the batch size and *τ* is the temperature parameter controlling the shape of the resulting distribution. The corresponding process is illustrated in Figure 2, where *x*_*i,t*_ is called the anchor, *x*_*i,t*+1_ the positive and *x*_*j,t*+1_, *j*≠ *i* are negative representations.

**Fig. 2:**
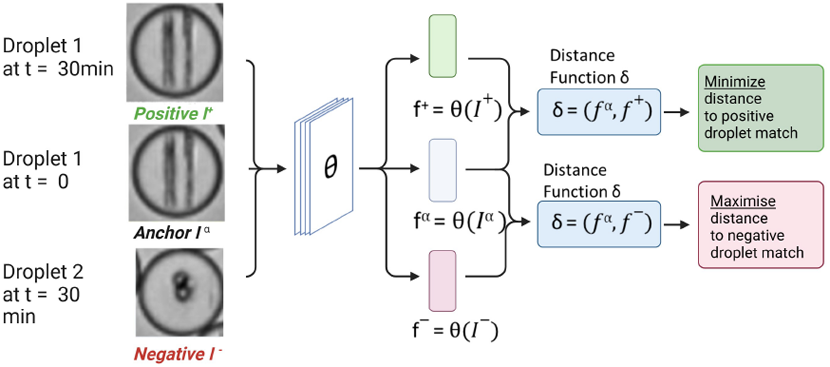
Contrastive learning of droplet representations labeled for Positive, Anchor and Negative as implemented in the algorithm Figure was adapted from [25].

**Fig. 3:**
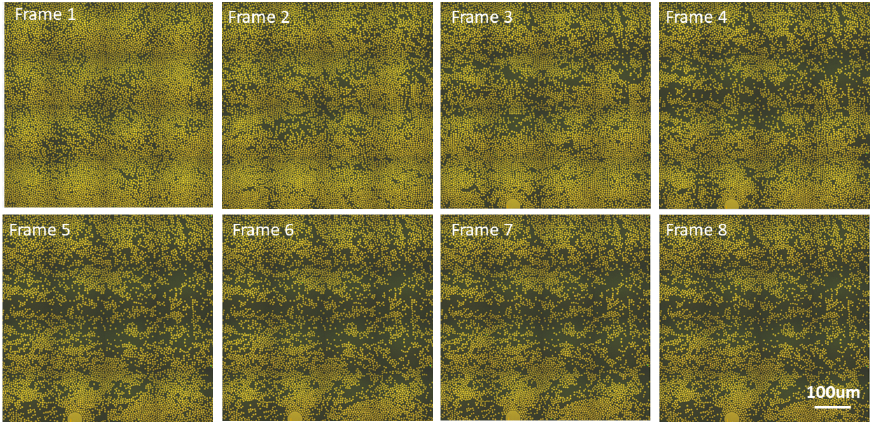
Visualization of example raw data from large-scale fluorescent micro-fluidic droplet experiments assayed at different time-points (frames). Varriyng degree of droplet movement (and thus cells contained inside) is visible between each frame increasing over-time.

### (E) Implementation of the OT algorithm for droplet tracking

In simple terms, OT tracking addresses the question of how to best transport a given source “mass” (droplet population) to a target mass (droplets in next frame) [23], [24]. We solved the actual OT problem, known as the fully unbalanced entropic regularized OT problem, using a slightly modified version of the commonly used Sinkhorn algorithm [26].

The input to OT is a cost matrix 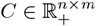. In our case *C* represents the cost of transporting a source droplet to a target droplet. The output of OT between two consecutive frames is a transport plan 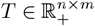, where *n* and *m* were the number of droplets in the source and target frames, respectively. The transport plan captures which droplets are supposed to be matched, with each entry *T*_*i,j*_ indicating how much mass is transported from the *i*-th droplet in the source to the *j*th droplet in the target frame. This results in the following optimization objective:

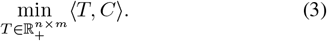

We designed a cost function that combined a positional loss and a visual embedding loss, according to:

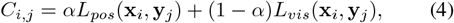

where *L*_*pos*_ and *L*_*vis*_ are the positional and visual embedding losses, *α* is a hyperparameter that determines the relative importance of each loss, and **x**_*i*_ and **y**_*j*_ represent the *i*^*th*^ droplet in the source frame and the *j*^*th*^ droplet in the target frame. More details on the encoder hyperparameter and metrics for encoder evaluation can be found in Appendix II.

Specifically, each loss was defined as follows:

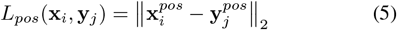

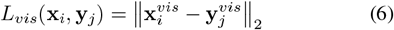

Further, we scaled the positional loss as such that the 95- quantile of the loss was numerically equivalent to the 95- quantile of the visual embeddings loss, making the interpolation between the two losses meaningful and controllable. In order to implement the basic OT function, two challenges had to be addressed. First, the Equation (4) had infeasible computational complexity, and second, the possibility of losing or gaining mass between frames had to be accounted for. To alleviate these two issues, the objective was modified to include entropy regularization for efficiency in Equation (3). Since droplets entered or left the frame, the number of droplets in the source and target frame might not be the same making the problem unbalanced^1^. Thus, Kullback-Leibler (KL) divergence terms were introduced to relax the balancedness constraints yielding the final objective as followed:

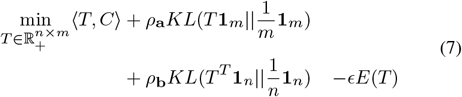

where **1**_*k*_ is the all ones vector of dimension *k*, while *ρ*_**a**_ and *ρ*_**b**_ control the amount of slack allowed in the balancedness of the problem. *ρ*_**a**_ controls balancedness of the source distribution and *ρ*_**b**_ the target distribution respectively, where high values enforce stricter control and balance of masses while lower values allow for slack in the conservation of masses. *E*(*T*) is defined as the entropy-regularizer of *T* ^2^, which controls smoothness and spread of transport matrices, encouraging more distributed or less sparse assignment of masses. Furthermore, the regularization parameter *ϵ* controlled the regularization strength and influenced how “willing” a given source point is to share its mass with multiple points of the target distribution rather than just transporting all its mass to a single point.

### (F). Trajectory generation

Lastly, we estimated trajectories for all droplets across all time steps using the solutions to the OT problem. The droplet detection mechanism assigned each droplet *i* in the source and *j* in the target mass unique IDs, 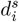 and 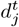, respectively. When a target droplet received the most mass from a given source droplet, we considered it as a match, *i*.*e*., we estimated that it was the identical droplet, and the target droplet received the same ID as the source droplet, which was later used to produce trajectories. Thus, the tracking assignment rule was defined as follows:

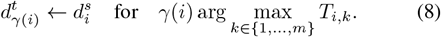

The transported mass, quantified by *s*_*i*_ = *T*_*i,γ*(*i*)_, indicates the certainty of the tracking of droplet *i* in the source to droplet *γ*(*i*) in the target frame. To increase the interpretability of the transport mass, we linearly scaled all entries in the OT matrix onto the range [0, 1] before applying Equation (8) to obtain 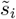, which provides an adequate approximation for confidence levels. To adjust 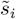 to represent true probabilities from which the confidence can be obtained directly, we calibrated 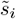 as described in the Appendix III. Using the matching rule (Equation 8) for all droplets and all frames of a time-lapse series, the trajectories of all droplets present in the first frame were predicted. If two source droplets were (theoretically) transported to the same target droplet, due to one droplet exiting the frame, the one droplet with lower amount of transported mass was removed and the corresponding trajectory filtered out.

### (G). Getting labeled data for CNN fine-tuning

In order to get data with high confidence we employed a heuristic approach by only considering a subset of droplets from an image array with little to no movement. Specifically, only droplets with less than 5 pixel of moved distance between frames were included as labeled data. This allowed us to extract droplets that were tracked correctly with a high likelihood. We validated this approach by manually checking the tracking results for a subset of droplets. This heuristic was used on a collection of 7 time-lapse series. We could extract 30k droplet pairs for training and another 30k droplet pairs for validation (the videos for training and validation were disjoint). To test the final model, we employed a data-bank of about 13k human-labeled pre-collected trajectories. This dataset is referred to as Fine-Tuning Data.

### (H). Movement, classification and time resolution

In order to independently evaluate the degree of total movement before tracking (from both experimental and simulated data), we calculated the Earth-Mover Distance (EMD). The EMD quantifies the “cost” of moving all objects between two time points by summing the total observed distance. To calculate the EMD, the input image is dimensionally reduced and treated as a single entity. The output EMD profile displays movement for each interval and median across the whole series. We calibrated the EMD metric against the actual distance (*µ*m) of average droplet movement in a separate image series, where a defined movement was introduced computationally VII. Within defined acquisition settings this allows translating the EMD to *µ*m distance or pixels moved. All images are classified based on their EMD/distance in to “low”,”low-to-medium”, “medium” and “large” movement as described in Figure 6. Initially, acquired experiments were taken at 1-2 min intervals to produce data that approximates labeled (ground truth) data, also referred to as “high-temporal resolution”. Such images require acquisition with small frame sizes (2-4 mm) in order to process the image in the fast interval time. Generally, unless movement is *>* 10 pixels (13 µm, per interval), this allows droplet tracking based on consecutive (droplet) centroid positions alone, which was validated using the droplet analysis software DropMap. From this data, subsets were generated by separating the image into lower temporal resolution series of 5 min, 15 min, and 30 min for direct comparison. These intervals mimick more common experimental settings usually obtained with larger dimension [18].

### (i). Measuring tracking failure using fluorescent droplet barcodes

Droplet experiments can contain fluorescent reporter probes for detecting secreted proteins. In validation experiments, reporter probes were replaced with fluorescent barcodes; these dyes encapsulated into each droplet, result in fixed, channel-specific intensities (barcodes) and can be generated using different ratios of dye to medium. Different dyes are encapsulated separately and mixed prior to acquisition. As background fluorescence (outside colored droplet) was typically 5-10,000 intensity units lower, the droplet intensity could be used as surrogates for correct trajectories. We developed Python code for reading out bio-assay or barcode intensities of droplets and cells by storing maximum, mean, and median fluorescent intensities. Taking generated trajectories (x,y per time-point) from out pipeline, we calculated the droplet fluorescent intensity around the estimated droplet position for each droplet,channel and time point respectively. Fluorescent intensity interchanges were calculated for each droplet time-transition (time point *T*_*x*_ compared to *T*_*x−*1_). An interchange was denoted “falsely tracked” if the intensity change was larger than 20% of the intensity distribution of all barcodes at *T*_0_. This metric then allowed to quantify the frequency of incorrectly tracked events per time interval.

## Results

In this work, we present an OT-based comprehensive pipeline to track objects from time-lapse imaging experiments. The degree and type of movement in droplet experiments can be large and highly diverse (Figure 3). Droplet populations can display areas of attraction, where droplets move towards each-other, repulsion, movement alongside an axis, as well as other complex and disordered motions. Because these movements are difficult to predict and the tracked objects are highly similar, we first investigated the contribution of different parameters, visual embeddings and other parts of the pipeline towards droplet tracking accuracy.

### (A). Validation of visual embeddings

First, we evaluated the visual embedding models on Dataset 1, containing magnetic nanoparticles (beadlines). In order to fine-tune the algorithm to discriminate between two different biological scenarios, two models were trained: Mod_cells_, which was trained exclusively on droplets with cells, and Mod_all_, which was trained on both droplets with and without cells.

We compared the effect of both models on the tracking accuracy from a validation set and a held-out test set 4. Overall, both models generated meaningful embeddings, allowing for effective droplet matching based on distances in the embedding space. Within the validation set, both models achieved a high top 1 accuracy of 0.952 ± 0.013 and 0.963 ± 0.011, indicating a high rate of correct matches for the nearest predicted droplet in the embedding space. In the smaller test set (test), they performed similarly well (0.983 ± 0.003 and 0.985 ± 0.02, respectively).

We noticed that increasing the sample size impacted the accuracy of matching based on distance only marginally, although the task became more difficult. The comparable performances on the validation and test sets suggested that the models were capable of generalizing across different time-lapse datasets. Interestingly, Mod_all_ slightly outperformed Mod_cells_ on top 1 and top 5 accuracy. As the performance was sufficiently high, we continued with Mod_all_ as the default pipeline.

To assess both the global and local structure of the embedding space, we reduced the embedding dimensionality using both UMAP (nearest neighbors=15, minimal distance=0.1, spread=10, Figure 4) and PCA (Supplemental Figure 5). For both dimensionality reduction techniques, we demonstrated that the same droplets, represented by the same-colored points, tended to form clusters according to their visual representation. Visual embedding models were capable of resolving single droplet trajectories based on visual features accurately, exemplified in 4B. While these evaluation datasets contained only minimal movement (*<* 50*µ*m average droplet movement), we concluded that OT-based matching should incorporate visual features for better performance.

**Fig. 4:**
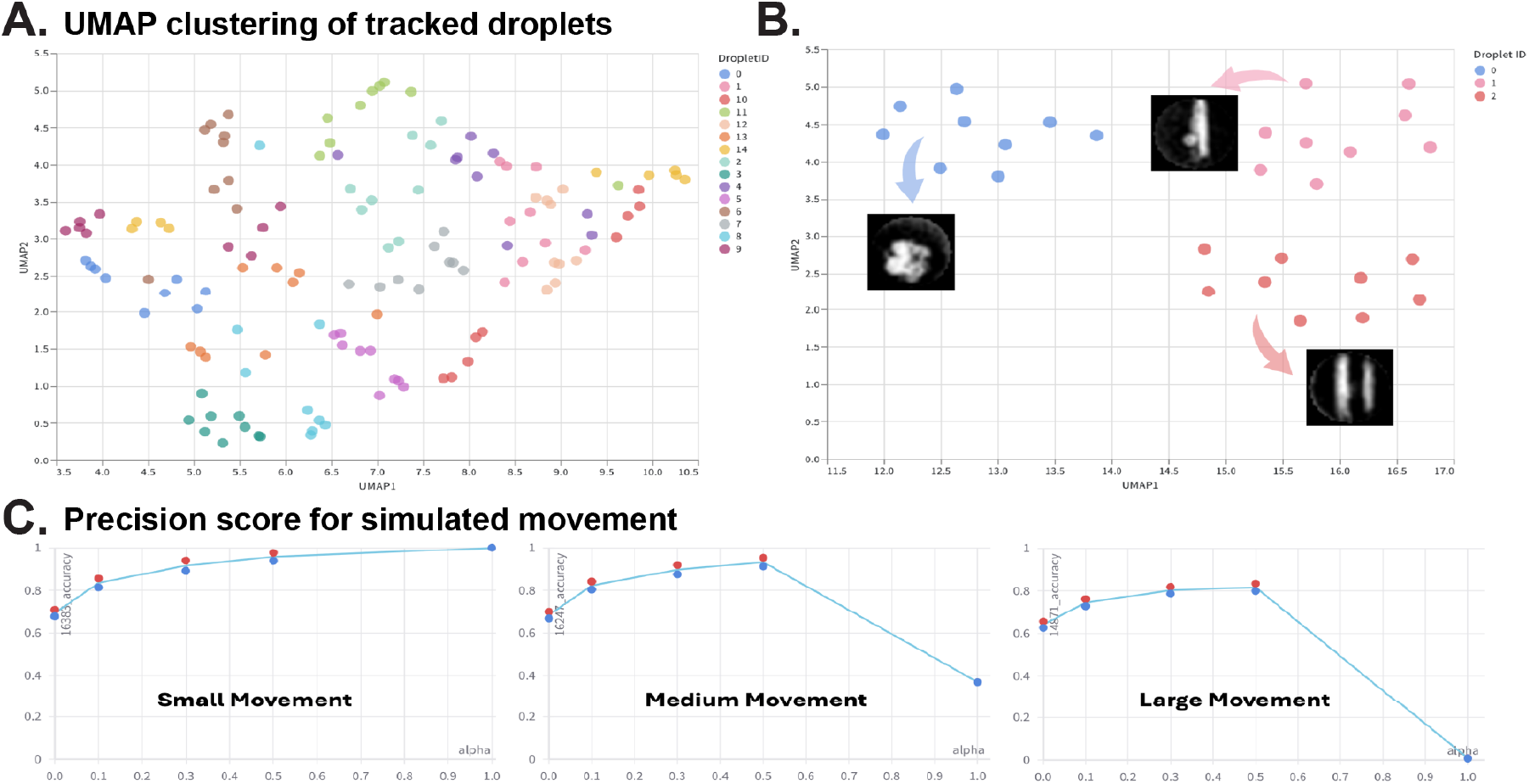
**(A.)** Embeddings of 15 random droplets tracked across 9 frames and reduced with UMAP to visualize tracking clustering performance, colored by their droplet ID. **(B.)** Embeddings of three representative droplets with different visual content were tracked across 9 frames. Droplet representations were reduced using UMAP to visualize tracking clustering performance, together with the original droplet images. **(C.)** Precision score for small, medium and large movement simulations as a function of *α*. Red and blue dots represent the runs with Model 1 and Model 2 used for the visual embeddings respectively. Other parameters: *ρ*_*a*_ = *ρ*_*b*_ = 0.999 for small and medium movement and *ρ*_*a*_ = *ρ*_*b*_ = 0.99 for large movement, *ϵ*_*rel*_ = 0.005. Embedding space cost defined through Euclidean distance. Corresponding Area under precision recall curve (AUPRC) scores for the best performing parameter sets for small, medium and large movement were 1.000, 0.998, 0.987 respectively.

### (B). Parameter search on simulated data

To quantitatively evaluate the full pipeline and to set the hyperparameters in Eq. 7, we performed a grid search on the regularization term in OT *ϵ*_*rel*_ (*ϵ* = *ϵ*_*rel*_mean(*C*)), on regularization parameters for adapting to data unbalancedness *ρ*_*a*_, *ρ*_*b*_ (set to either 0.99 or 0.999), and on the weighting parameter which controls the contribution of location loss to the cost function *α*. All results were generated using the simulated data, which can be found alongside tested values in Appendix IV-C. Our simulation had 20k droplets in each frame, but we cut image boundaries in silico to introduce (more realistic) imbalance, leaving about 16k droplets with some degree of imbalance across different frames.

We observed that irrespective of other parameters, the choice of *ϵ*_*rel*_ did not significantly influence the metrics. Thus, we choose *ϵ*_*rel*_ = 0.005. The choice of *ρ*_*a*_ = *ρ*_*b*_ = 0.999 was optimal for small and medium movement data while *ρ*_*a*_ = *ρ*_*b*_ = 0.99 was better for large movement. Hence, we plotted the metrics according to this choice and subsequent runs were obtained with these selected parameters.

Finally, we studied the precision for all droplets and of the top 10,000 most certain droplet trajectories (precision@10k). We clearly observed that bigger values of *α* worked better for smaller movement, while for medium and large movement taking an alpha of about 0.5 performed best (Figure 4, Supplemental Figure 4). We concluded that for small movement (EMD *<* 0.5), just using the positions was almost sufficient for the OT to work efficiently and as soon as there is more movement we had to include visual information to get high scores. As biological conclusions require precision on droplet populations, false negatives are preferred over false positives. Hence, droplet populations successfully tracked should be correct with high confidence even when filtering out droplets that can be potentially tracked. Therefore, for the intended use of the algorithm where trajectories with high certainty are provided, we recommend using *α* = 0.5, which yielded almost perfect tracking for 10k most certain droplets in our experiment, which is more than 50% of the droplets in scope.

### (C). Visual evaluation of reported trajectories

We next visualized and validated the droplet trajectories returned by our method. Figure 5 presents an analysis of a small section of the large droplet array. The predicted droplet centers are shown as interconnected dots overlaid on the real microscope image from the first step, and the lines provide the trajectory of the center of the droplet.

**Fig. 5:**
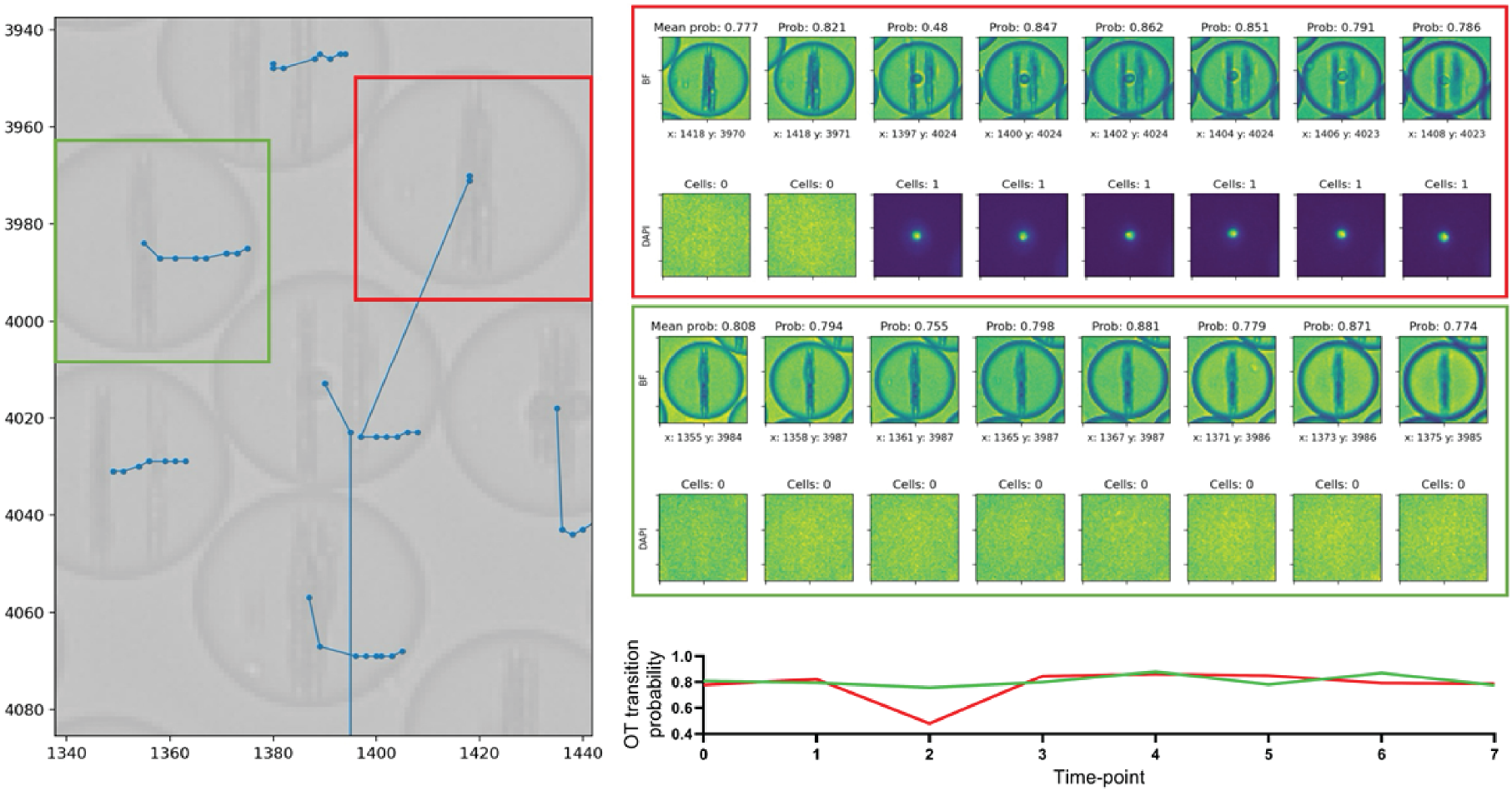
Overlay of droplet regions with tracking positions from initial tracking analysis of small-movement experiments. Blue dots and lines indicate trajectories over all eight time frames. The background shows the bright-field channel of the first frame. Green bounding box indicates high-confidence tracked droplet across time intervals. Red indicates a mismatch. Corresponding, extracted Bright-field and DAPI channel patches of two droplet trajectories are depicted with Optimal transport (OT) probabilities for each time transition. Indicated OT transition probabilities (Prob.) refer to each transition from previous position. The mean probability of the whole trajectory is included above the first time frame (Mean prob.). Information about the detected number of cells is included above the DAPI channel at each time frame as OT transition probability graphed.

While some trajectories appeared plausible, such as the one from the droplet in the green box, we could identify mismatches (indicated by the red box) visually and based on OT-scores between the second to the third frame. In Figure 5, the green and red isolated droplet patches were displayed alongside transition probabilities for the two predicted trajectories. This discrepancy was evident also in the OT transition probability, which dropped to 0.48 during this transition but otherwise stayed above 0.75, as shown by the line-plot below. We concluded that observed irregularities (red series) could also be identified in the post-tracking analysis visually or by the OT-transition score.

### D. OT-based tracking produces high accuracy and more than 80% increase in trajectory recovery

On the High-Temporal-Resolution data, we ran the best configuration of our pipeline, as determined during parameter search. We applied the same settings to 5 min, 15 min, and 30 min resolution images and calculated what percentage of trajectories could be recovered compared to high-temporal resolution. As a comparison or baseline frequency, we used the percentage of droplets, which can be tracked using the Matlab-based data analysis software (DropMap), [5], which allows tracking of movement *<* 10 pixels. We showed that only 23% of droplets in low-movement images can be tracked and analyzed using this method, while images with larger movement resulted in complete loss of data (Figure 6). To compare experiments in terms of total image movement (small, medium, and large), we calculated the Earth-Mover distance metric (EMD) for each experiment. The small movement exhibited a median EMD of 0.483, medium – 0.895, and large images – up to 6.294 (Supplemental Figure 8). We converted the EMD to the average distance moved in *µ*m and droplet diameter to quantify the droplet movement (Supplemental Figure 10). Small movement images demonstrated an average movement of all droplets of 2 diameters, medium of 12 diameters and large movement of up to 160 diameters, corresponding to 10.000 micrometers.

**Fig. 6:**
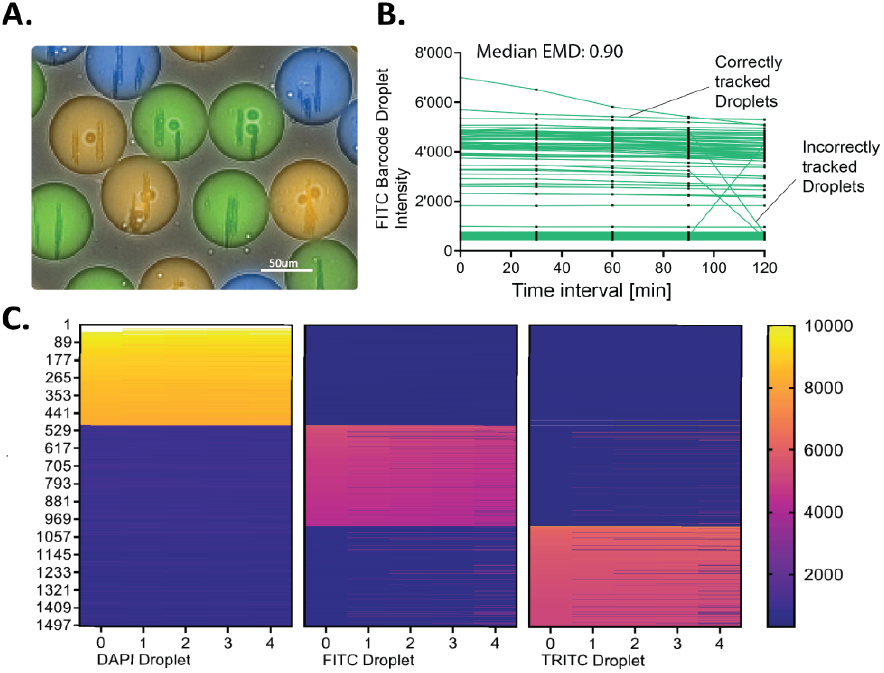
**A**. Representative droplet image of barcoded experiments. **B**. FITC (Green) Mean fluorescence intensity (MFI) of 500 droplets picked random over each time-point for experiments with human PBMCs and beadline bio-assay from medium movement 30-minute time intervals. Highlighted are example correctly and incorrectly classified trajectories according to intensity change between intervals. **C**. Heatmap of individual droplet trajectories of 500 droplet sorted by channel and intensity in descending order. X-axis represents time-points (5 time-points) from 30-min interval experiments. Colors indicate fluorescence intensity.

In terms of image tracking accuracy, we observed a downward trend in accuracy for trajectories as time resolution decreased (Figure 6). Nevertheless, we maintained an overall accuracy greater than 0.6 even at 30-minute intervals and for large movements. The downward trend makes intuitive sense, as there was more unobserved information between consecutive frames. To put in perspective, we state the tracking success of the 30-minute intervals as the frequency of droplet OT-scores which are above 0.6 and above 0.77, in alignment with figure 5 (The average OT-score for trajectories with 1 false droplet in 30-minute intervals ranges between 0.4 and 0.75). The frequency of droplets where any transition between timepoints is above 0.5 OT-scores is 91.6, 99.3 and 61.8% for the different movement types respectively. Thus the results show, that overall droplet tracking was improved from previously 0% to over 90% in some instances. This demonstrates the strength of the pipeline in recovering trajectory data. A Cutoff of 0.5 is chosen according to the OT-score expected, when a trajectory resulted in a false transition (Figure 5). To note, the discrepancies in tracked events between medium and large movement (in terms of 0.7% tracked events versus EMD) is due to a small sub-populations of droplets moving little (¡ 10 pixels), while the majority moves larger distances. It is important to recognize that movement may not be a uniform, normally distributed phenomenon, but rather incoherent, with some cells and droplets moving large distances while other can remain still. We conclude that even images with EMD *>* 2 can be accurately tracked terms of correct trajectories.

### E. Confirmation of tracking success using fluorescence intensity barcodes

To externally validate the resolution at which droplet trajectory can be recovered from time-lapse series, we added fluorescent barcodes into each droplet, consisting of fluorescent-colored dyes, for example, FITC (Green) combined with cell staining in DAPI (Blue). The fluorescence was not used for tracking, but the constant, channel-specific intensity allowed to identify non-correctly tracked droplets in a simple manner. In the case of an incorrect matching, the centroid (x,y)-position would result in a large spike or fall in intensity (e.g. FITC-to-TRITC or FITC-to-background). This interchange was calculated for every droplet-time transition and dependent on the intensity change (increase or decrease) of each droplet at timepoint *T*_*x*_ compared to *T*_*x−*1_.

First, we investigated whether we could reconstruct trajectories using barcode intensities. Fluorescent barcodes could reproduce excellent tracking qualities and accuracies in accordance with previous data (Figure 6). For medium movement, the tracking of images with trained machine-learning approach resulted in 99% accuracy in droplet tracking with results comparable to high-temporal resolution (Figure 6A), supporting OT-score data. Visual representation of the FITC channel intensities over time (Figure 6B) demonstrated that only very few, distinct intensity interchanges were observed.

Finally, calculated trajectories could also be visualized in a heat-map format where droplets were mapped according to channel (barcode) and sorted according to intensity (Figure 6C). Each line in the heatmap indicated a droplet tracked over the whole time course of the experiment. Particular noteworthy is that tracking of droplets containing cells (DAPI, left-hand), almost no transitions were observed.

### F. With cells only, high-accuracy tracking can be maintained for a sub-population of droplets

Finally, we investigated if droplets with altered visual features and droplet content can be tracked with high resolution without fine-tuning the algorithm. For this purpose, we implemented dataset 3, consisting of droplets containing one or multiple CHO cells, but no nanoparticles. With one or multiple unknown cells, potentially interacting with eachother inside droplets, to produce high-resolution, single-cell data from these images demonstrate a typical experimental and data analysis challenge (Figure 7A). Applying the algorithm to these images led to worsening of tracking qualities, in particular for large movements in line with previous results (Figure 7B). For low-to-medium movement, incorrectly assigned barcodes occurred in around 30% of droplets at 200 minutes of tracking and in 50% of droplets at the last interval (Figure 7C). However, overall, approximately 30-40% of droplets could still be tracked without any false transitions. Whereas not ideal, this still demonstrated success compared to previous algorithms, where low movement images could be tracked at around 20%. Loss of tracked droplets in the “low movement” condition (median EMD: 0.25) can also be attributed to movement late in the time series as indicated in the EMD profile (Supplemental F igure 9). Interestingly, cells (labeled in blue, DAPI) could again be tracked with high accuracy within the same experiment across all frames. The higher precision of cell-containing droplets is also apparent in the heatmap: Droplets containing cells show correct and stable trajectories of the fluorescent barcode with much higher frequency compared to barcodes alone (Figure 7B).

**Fig. 7:**
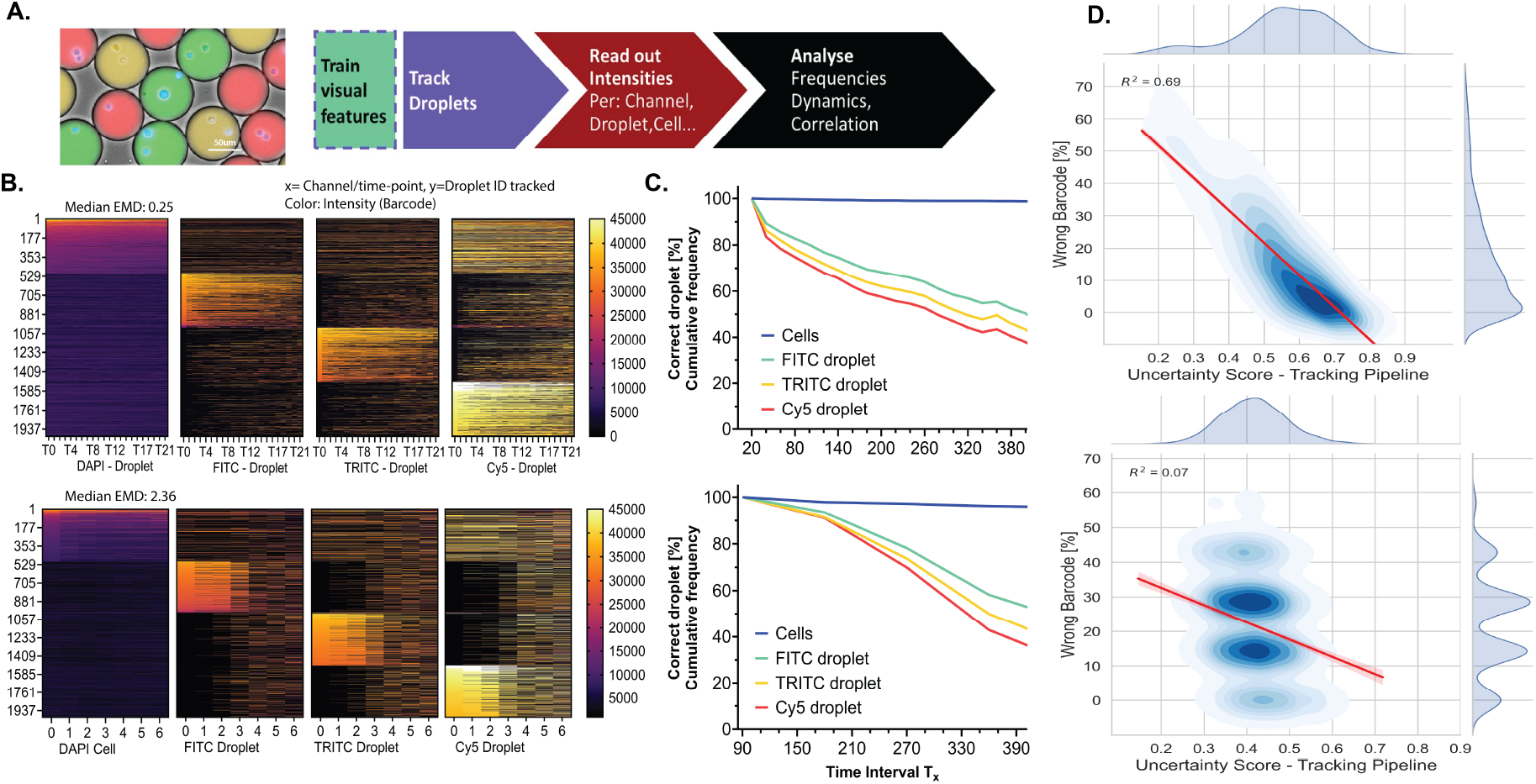
CHO cell tracking and barcode analysis. Read-out of droplet average intensity and cell signal from generated trajectories. **A**. Exemplary image of barcoded droplets with three droplet fluorescence dyes and DAPI-violet stained cells. Workflow for the training, tracking, read-out and analysis of biological in-droplet experiments. **B-D**. Analysis of two independent experiments with color-barcoded droplets and CHO cells from experiments without additional visual features (beadline) and without prior data training, with low-to-medium movement (EMD: 0.249, upper panel) and medium-to-large movement (EMD: 2.36, lower panel). **B**. High resolution data visualization of barcode intensities (color) of individual droplet trajectories from left *t*_0_ to right *t*_final_. Per channel 500 droplets were picked at random and sorted according to fluorescence intensity in descending order for better visualization. Each line in the heat map represents a tracked droplet trajectory ID (y) over time (x). **C**. Cumulative frequency of successful tracking separated for each barcode channel as well as for droplets containing cells (DAPI: Violet) for both movement-type images. **D**. Correlation between the mean uncertainty OT-score of each droplet and the percent instances that a wrong color was identified from bar-coding. The red line indicates the linear regression curve and R-squared value for the data correlation.

For large movement images (Figure 7C), failure to track occurred in 20% of the events after 200 minutes of measurement and 40-50% at the end of the measurement, stemming from additional (challenging) movement late in time-lapse series (Figure 7B, Supplemental Figure 9). The frequency of droplets with perfect trajectories in large-movement images was 30% at the end of the measurement. As this was only marginally weaker compared to low-to-medium movement, it indicated that movement alone did not solely predict whether the algorithm could produce accurate trajectories, but the accuracy was rather attributed to a combination of visual content, image resolution, and optimal transport. Finally, the correlation of the uncertainty score produced by the pipeline and the occurrence of a false transition (Figure 7D, linear regression depicted as red line with R2 value) showed that the trajectory uncertainty score (output of the tracking model) is a good predictor and surrogate for filtering “correct” droplets. Droplets with higher uncertainty scores showed lower transitions, in line with Figure 5. Overall, we concluded that OT tracking was able to recover substantial trajectories for low-movement conditions even in more challenging experiments. When large movement is present OT-score filtering may still permit to retain a tracked subpopulations with correct trajectories.

## Conclusion and Discussion

We present a general workflow for the data-independent tracking of circular objects (droplets) and cells from large scale over-time microscopic images. The described platform is the first p roof-of-concept a lgorithm t hat u ses unsupervised tracking of droplets containing cell and immuno-assays over time and across space by using the theory of optimal transport coupled with visual feature training. The implementation of solving the OT problem for complex and disordered movement in fluidics systems of droplets provides an end-to-end pipeline for analysis.

Our results suggest that whereas classical machine-learning (visual feature training and validation), together with positional data, is sufficient for low movements, more comprehensive algorithms are needed for images with larger movements (EMD *>* 0.5) and longer time intervals. Our analysis of attraction and repulsion, along with the optimization of visual features and integration of droplet features with positional data, resulted in a comprehensive model. Investigating newly generated trajectories using low-dimensional clustering resulted in clustering of visually similar droplets from a heterogeneous population. Moreover, the success in difficultto-track images and long time-interval instances (30 minutes) from previously 0-0.7% tracked droplets to 60-99.3% can be heavily attributed to the precise configuration and multifactor integration of the pipeline. Barcode readout mimicking our biological assay confirmed these results and enabled the construction of complete and correct trajectories for DAPI (Cell), FITC, TRITC, and Cy5) fluorescence intensities over time for different degree of movement. When challenging the pipeline with novel visual features (and the absence of nanoparticles) as in the CHO cell experiments, we could still maintain tracking for 40% in low and for 30% of droplets in high movement conditions throughout the time series.

Quantitative analysis of our tracking yielded metrics such as the OT score, the accuracy of which could be confirmed as well using droplet barcodes. This metric can be used for a reliable criterion to select successfully tracked droplets. In fact, using the OT-score on our CHO cell dataset yielded a subset of droplets which can be tracked and filtered out with high certainty, demonstrating advancement also at the analysis stage. Finally, successful tracking of DAPI-stained cells across different experiments compared to droplets without cells indicates that visual features (cells inside droplets) still contribute significantly to the pipeline. We conclude that while optimal transport enables more comprehensive tracking at the population level, visual features may require re-training or calibration in instances where droplet size or cell morphology is altered and/or if other visual features (such as nanoparticles) are not present. In this context, it is important to emphasize the importance of balancing parameters between OT and visual features, as well as the role of labeling. In particular, for large movements, the pipeline requires visual data inside each droplet, which is detrimental to high precision. Hence, some visual labeling may still be important when aiming to track completely novel droplet content in long-duration or high-movement experiments. As broadness in analysis parameters came at some cost in precision for cell and droplet tracking when tested on novel experiments, in the future, refining tracking models may result in even better adaptability across datasets. In fact, evaluation of movement metrics (EMD), droplet tracking quality (OT-score) and overall image quality could enable potentially automated fine-tuning o f t he algorithm. Such fine-tuning c ould i nclude i ncorporating training to optimize the visual feature space and to set and optimize the balance between OT and visual features. Overall, we believe our approach has biological applicability, being able to retrieve correct droplets, enabling more precise analysis with metrics for quantity and quality of produced trajectories and increasing throughput. Therefore, with fine-tuning parameters, our workflow could also be applied to other areas, such as for drug-organoid screening or non-droplet applications, including the tracking of extracellular vesicles or data from life-cell imaging [28] [29].

## Author contributions

VB, IL and KE designed the study and supervised the project. MV, SG, TH, RD, and WO constructed the tracking pipeline workflow and simulations. AL and IL generated initial cytokine detection tracking experiments. LS generated cell experiments and constructed droplet read-out and analysis workflow. L S a nd M V w rote t he i nitial d raft, a nd a ll authors commented and revised the manuscript.

## Acknowledgments

We want to acknowledge the initial algorithms for droplet, cell detection, pre-processing and visualization which were implemented by Data Science Students from the ETH AI Center Data Science Lab 2022, namely Antoine Basseto, Fransceco Da Dalt, Victor Gillioz, Samyak Jain, Filipe Cunha, Haruki Shirakami, and Yihao Liu.

## Appendix I Preprocessing

### A. Droplet Detection

We detected droplets in the BF channel. For this, we first applied rank equalization, to have more contrast in the image and then a Gaussian blur filter to remove noise from the images. The detection of droplets in the image is performed using the Hough Circles algorithm, from the OpenCV library [19]. The algorithm is based on a three-parameter description of a circle, first to detect edges in the input image and then, given a specific predefined radius, to find the circle center. In our case each droplet is a circle. We look for radii of different sizes, to capture all droplets. Then all found droplet parameters (center and radius) are refined using the RANSAC (Random sample consensus) algorithm to estimate model parameters in the presence of outliers [20].

#### B. Visual Embeddings

For the preprocessing of the raw data for visual embeddings, we process the BF and DAPI channel differently: For the BF channel, we first re-scaled the intensity to [0, 1] and then applied median blurring and rank equalization. Then we set all values which are below the 50%-percentile of the intensity values to zero. For the DAPI channel we set all values which are below the 80%-percentile of the intensity to zero and then apply the same steps as in the BF channel. Supplemental Figure 1 shows the output of pre-processing, a detected droplet (raw data), droplet and cell after pre-processing for droplet selection and feature extraction for comparison. Extracted features are labeled (manually) in red.

**Supplementary Figure 1:**
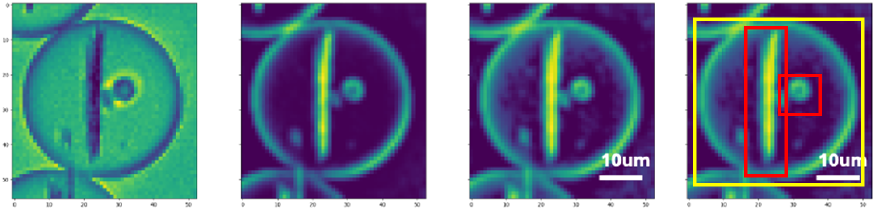
Droplet extraction during and after pre-processing. **(Left)** Raw image of cropped droplet. **(Middle)** Preprocessed for droplet detection. **(Right)** Preprocessed for feature extraction. Extracted features are overlayed from bright-field channel and labeled (manually) in red. Detected droplet marked in yellow.

#### C. Cell Detection

After data preprocessing for droplet detection, we can also use the detected droplet for counting the number of cells in each droplet. We cut out each droplet in the preprocessed DAPI channel to detect whether cells are present in the given droplet or not. To find a c ell, w e fi rst us e th e NM S (nonmaximum suppression) algorithm to get an initial guess of where cells could be in a droplet. Then, using a threshold, we check whether the persistency and intensity at the proposed position are high enough to be classified a s a c ell a nd count the number of cells per droplet. We note that the algorithm for droplet detection is not guaranteed to detect all droplets in the image. Similarly, cell detection may fail to recognize weakly stained cells or struggle in situations with low resolution or blurred data. However, from manual, visual inspection and displaying of droplets, we we are confident that most if not all droplets are detected. We use this detection as a ground truth acceptance in the subsequent model.

## Appendix II

### Encoder Training and Visual Embedding Metrics

#### A. Choosing the encoder hyperparameter

To gauge the quality of the visual embeddings, we mainly use the accuracy of matched droplets across two frames. We compare each droplet with its 128 nearest neighbors in the next frame and consider the droplet to be correctly matched if the distance to its positive match is minimal among all distances computed. Further, we consider two different sets of training data; (i) we consider droplets which contain cells only and (ii) to use all droplets.

**Supplementary Table 1:** Encoder hyperparameter settings for training the encoder. Learning Rate, Batch Size and Embedding Dimension

|  |  |
| --- | --- |
| Optimizer | Adam [30] |
| Learning Rate | 0.001 |
| Batch Size | 128 |
| Embedding Dimension | 20 |

##### B. Metrics for evaluating the encoder

For evaluating the embeddings we consider two metrics: accuracy and area under the ROC curve. We report the obtained values of the metrics in Section III-B, Table II.

**TABLE I:** Overview of experiments and generated datasets, their biological application, and what they were used for in tracking analysis. Exp. Experimental, Sim. Simulated.

| Datasets | Type | Droplet content | Application | Used for: |
| --- | --- | --- | --- | --- |
| Dataset 1 | Exp. | Beadline + PBMCs | Cytokine secretion | Biological validation and tuning |
| Dataset 2 | Exp. | CHO cells | Cell reporter | Cross-validation |
| Dataset 3 | Sim. | Beadline and cells | Simulating (modeling) movement | Labeling validation and accuracy analysis |

**TABLE II:** Performance metrics for different models across validation and test datasets. Top 1 and top 5 accuracy as well as area under the ROC curve (mean and its standard error across 9 frames). Number of droplets in the test dataset: 13584. Metrics are based on Euclidean distance. A detailed description of the metrics can be found in the Appendix II. Val., validation.

| Metric | $\text{Mod}_{\text{cells}}$<br>(val.) | $\text{Mod}_{\text{cells}}$<br>(test-sub.) | $\text{Mod}_{\text{cells}}$<br>(test) | $\text{Mod}_{\text{all}}$<br>(val.) | $\text{Mod}_{\text{all}}$<br>(test-sub.) | $\text{Mod}_{\text{all}}$<br>(test) |
| --- | --- | --- | --- | --- | --- | --- |
| TOP 1<br>ACC | 0.952<br>(0.013) | 0.992<br>(0.002) | 0.983<br>(0.003) | 0.963<br>(0.011) | 0.993<br>(0.001) | 0.985<br>(0.002) |
| TOP 5<br>ACC | 0.981<br>(0.008) | 0.999<br>(0.001) | 0.996<br>(0.001) | 0.986<br>(0.006) | 0.999<br>(0.000) | 0.997<br>(0.001) |
| AUROC | 0.988<br>(0.001) | 0.990<br>(0.000) | 0.990<br>(0.000) | 0.988<br>(0.001) | 0.990<br>(0.000) | 0.990<br>(0.000) |

- **Top 1 and top 5 accuracy:** We assumed a dataset of matched droplets. For a given droplet embedding we compute its k-nearest neighbors from embeddings of all droplets of the consecutive frame. If the matched droplet embedding is present in the k-nearest neighbors we considered it a positive match. The accuracy is the ratio of positive matches to the number of considered droplets.
- **Area under the ROC curve:** To obtain the ROC curve, we computed the distances between embeddings of correctly matched droplets (positive samples) and between randomly matched droplets (negative samples). Then we concatenated the two lists of distances and sort them in increasing order. For a given *k* we observed the first *k* elements and call the fraction of the ones coming from positive samples a true-positive rate (TPR) and the fraction of the ones coming from the negative samples a false-positive rate (FPR). By varying *k* we obtained paired values of TPRs, and FPRs which we plotted as the ROC curve. Finally, we computed the area under the curve.

Visual embeddings can be accessed online: (Interactive embedding visualization), where by hovering with the mouse cursor over the point one can see an underlying droplet image. From this visualization, we observed that droplets with similar magnetic bids (their number and width) tend to be located close to each other in the reduced embedding space.

## Appendix III Model

### Calibration

Calibration curves, also known as reliability diagrams, provide a visual assessment of the calibration of probabilistic predictions from a binary classifier. These curves plot the predicted probability against the positive label frequency, offering insights into how well the model’s predictions align with actual outcomes (Examples shown in Supplemental Figure 2). We employed isotonic regression to map the match scores 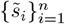 to calibrated probabilities 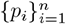, in order to align the predicted scores with the observed frequencies of positive outcomes. Given a set of predicted scores 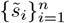 and the corresponding observed binary ground truth 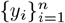, isotonic regression finds a monotonic, non-decreasing *f* : ℝ → [0, 1] which minimizes the following objective:

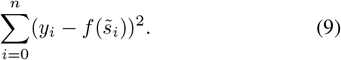

The calibrated probabilities are then given by 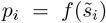 for *i* = 1, …, *n*. The monotonicity constraint ensures that the recalibrated probabilities maintain their order, preserving the relative ranking of samples.

**Supplementary Figure 2:**
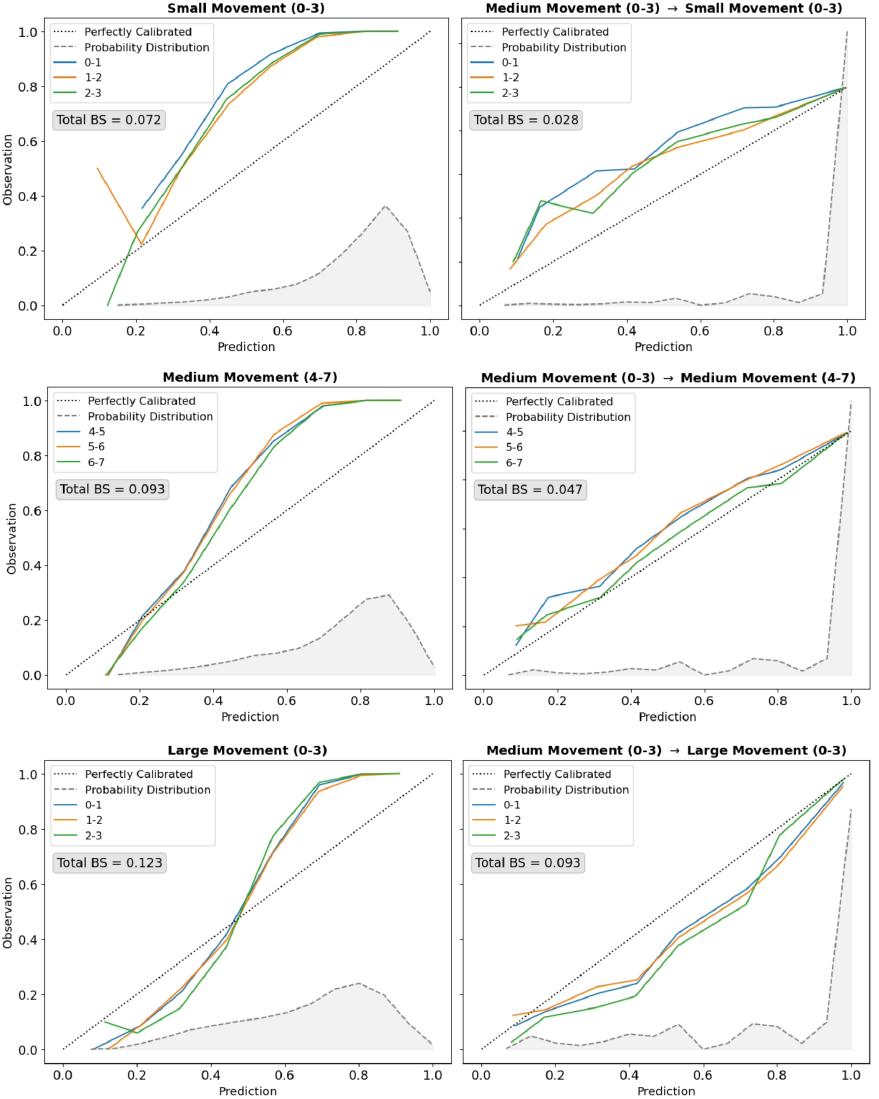
**(Top)** Model calibration for small movement, trained on medium movement. **(Middle)** Model calibration for medium movement, trained on medium movement. **(Bottom)** Model calibration for large movement, trained on medium movement.

In scikit-learn algorithm, the isotonic regression solver is based on the Pool-Adjacent-Violators Algorithm (PAVA) [31] [32]. In the context of our pipeline, we train the isotonic regression model on simulated data and then apply it when performing tracking on other real or simulated data.

**Supplementary Figure 3:**
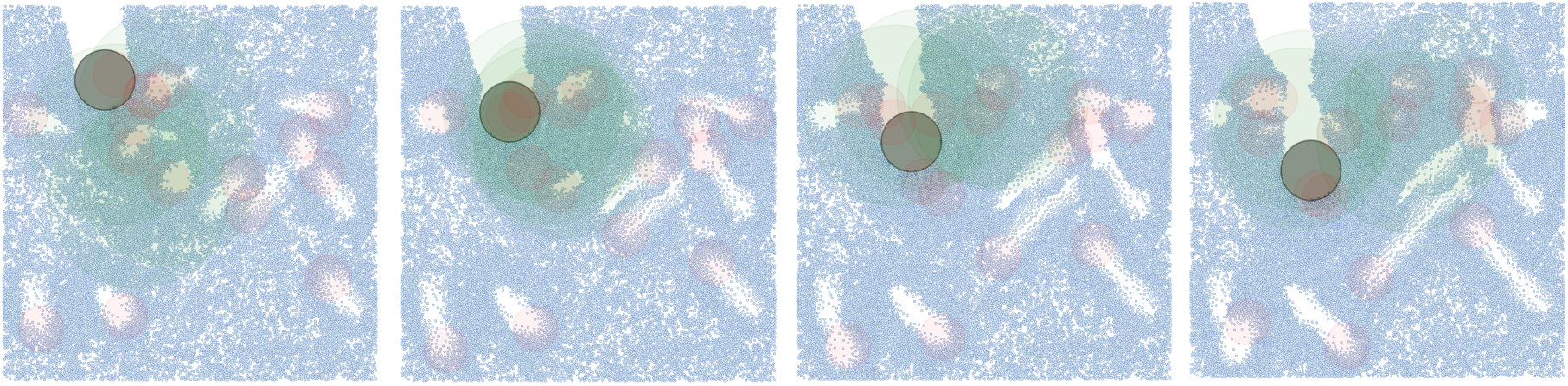
Simulation of positions of 20k droplets across a time-lapse series of n=4 from left to right, for medium movement. Model shows irregular movement, traversing the image as well as areas of attraction and repulsion.

Supplemental Figure 2 shows calibration plots for the tracking of around 8k droplets for small, medium and large movement. The title indicates the type of movement and the frame range on which we applied our evaluation pipeline. For all figures, calibration plots for which we used the raw scaled OT matrix entries as confidence levels for the match predictions are displayed on the left. On the right, one can see the calibration plots for the same trajectories on the same type of movement but with calibrated probabilities. Here we calibrated by training the isotonic regression model on the results for medium movement across frames *F*_0_ to *F*_3_ and applying it to the respective match scores. To avoid testing on training data, for the calibration plot in Supplemental Figure 2b different frames of the simulation and different image patches were used to generate the results. The dotted line indicates a perfectly calibrated model, the area marked in grey shows the distribution of predicted confidence scores and the Brier score across all frames is given in the grey box (lower values are desirable). Differently colored lines correspond to the predictions for transitions between different pairs of frames.

## Appendix IV

### Simulated data

#### A. Creation of labeled data set

Applying our method to real data with very little movement, we were able to predict a set of trajectories for which we are confident that the majority thereof are true positives. This confidence stems from visual observations using the visualizer tool and filtering based on the assigned uncertainty from the algorithm, distance from the image segment border (at least 20 pixels) and amount of movement (at most 5 pixels).

We then combine the matched patches with simulated position data to imitate the real data with significant movement. This allows us to quantitatively evaluate our method on various kinds of movement, while still relying on visual embeddings.

#### B. Pipeline on Simulated Data

The pipeline for processing simulated data accepts as input a .csv file with obtained droplet locations as well as a .npy file with matched droplet patches and a .csv with the number of cells per droplets. Instead of providing the last two files, one can obtain them using the pipeline itself. This requires running our code on the real low-movement image first, to obtain some high-confidence trajectories. The pipeline matches provided location data with the patches and the number of cells, keeping track of the droplet ID across frames. The next steps, until obtaining calibrated trajectories are identical to running the pipeline on the real data. Since the pipeline keeps track of the droplet ID across frames, at the end of the run, metrics for the trajectories are computed and reported.

#### C. Parameter search for simulated data

In order to find optimal parameters for our pipeline we have run an extensive parameter search. Due to the task being computationally heavy, we first run a finer search on the simulated data with 6000 droplets across 4 frames and after selecting promising subset of parameters we rerun the experiment on data with 20000 droplets across 4 frames and repeat all the experiments for small, medium and large movement simulation. The parameters we consider were:

- *α* ∈ {0, 0.1, 0.3, 0.5, 1} which represents the contribution of the location loss to the cost function. *α* = 0 indicates scenarios when using only positional information and *α* = 1 for only visual information.
- *ρ*_*a*_ = *ρ*_*b*_ ∈ {0.99, 0.999} which requires adaptation to the unbalancedness of the data.
- *ϵ*_*rel*_ ∈ {5 · 10^*−*3^, 5 ·10^*−*4^, 5 ·10^*−*5^} which specifies the regularising *ϵ* in the OT as *ϵ*_*rel*_ times the mean entry of the cost matrix.
- Visual embedding model which can either be Model_cell_ trained only on droplets with cells or Model_all_ trained on both droplets with and without cells.

#### D. Simulation Evaluation Metrics

The advantage of having simulated ground-truth data is that we can assess the quality of tracking quantitatively without expert labeling. This allows for tuning of OT hyper parameters and choosing a suitable contrastive learning model. For every predicted matching of droplet *i* and some other droplet between a source and target frame, let *p*_*i*_ ∈ [0, 1] be the assigned confidence level and *y*_*i*_ ∈ {0, 1} indicate whether the match is a true or false positive. Then several metrics can be computed using the set of tuples 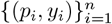, where *n* is the number of droplets in the source frame. We can also compute any of these across all frames to obtain total metrics describing the performance on whole trajectories.

#### E. Precision

A simple way to evaluate the performance of our tracking approach is to assess the precision given as:

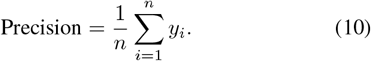

*1) 10K precision:* The precision within the top *k* choices of the tracking method. Let *r*_*i*_ = rank(*i*) be the rank of the *i*^*th*^ droplet ordered in descending order with respect to *p*_*i*_ (i.e., highest to lowest probability). Then the Precision@k is given as:

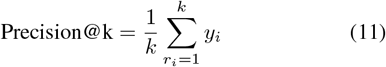

The precision for 10k droplets is presented in Supplemental Figure 4.

## Appendix V

### Visualization

In order to visually evaluate the performance of our method on real datasets two visualizers for trajectory curves were implemented into the pipeline, originally designed by ETH Data Science Lab (DSL). I. Droplet position visualizer and II. a visualizer for matched, extracted droplets across time series and channels. Visualization was used, for instance in Figure 5.

### Appendix VI

#### PCA versus UMP analysis of batch effect in droplet classification

**Supplementary Figure 4:**
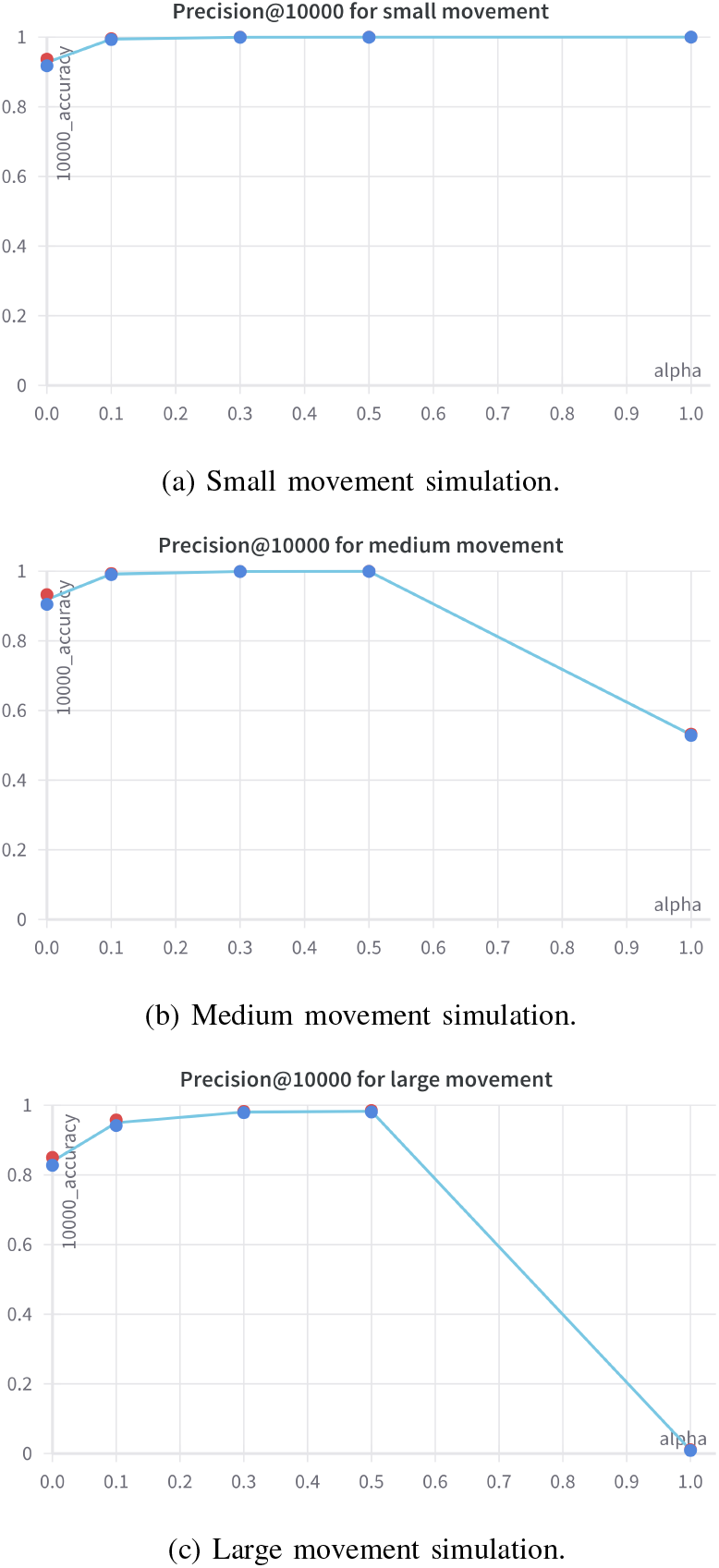
Precision score for small **(a)**, medium **(b)**, and large **(c)** movement x simulations for 10K droplets as a function of *α*. Red and blue dots represent runs with Model 1 and Model 2 used for the visual embeddings, respectively. Other parameters are configured as: *ρ*_*a*_ = *ρ*_*b*_ = 0.999 for small and medium movement, and *ρ*_*a*_ = *ρ*_*b*_ = 0.99 for large movement, *ϵ*_*rel*_ = 0.005. The embedding space cost is defined through the Euclidean distance. Corresponding AUPRC scores for the best performing parameter sets for small, medium, and large movement were 1.000, 0.998, and 0.987, respectively.

**Supplementary Figure 5:**
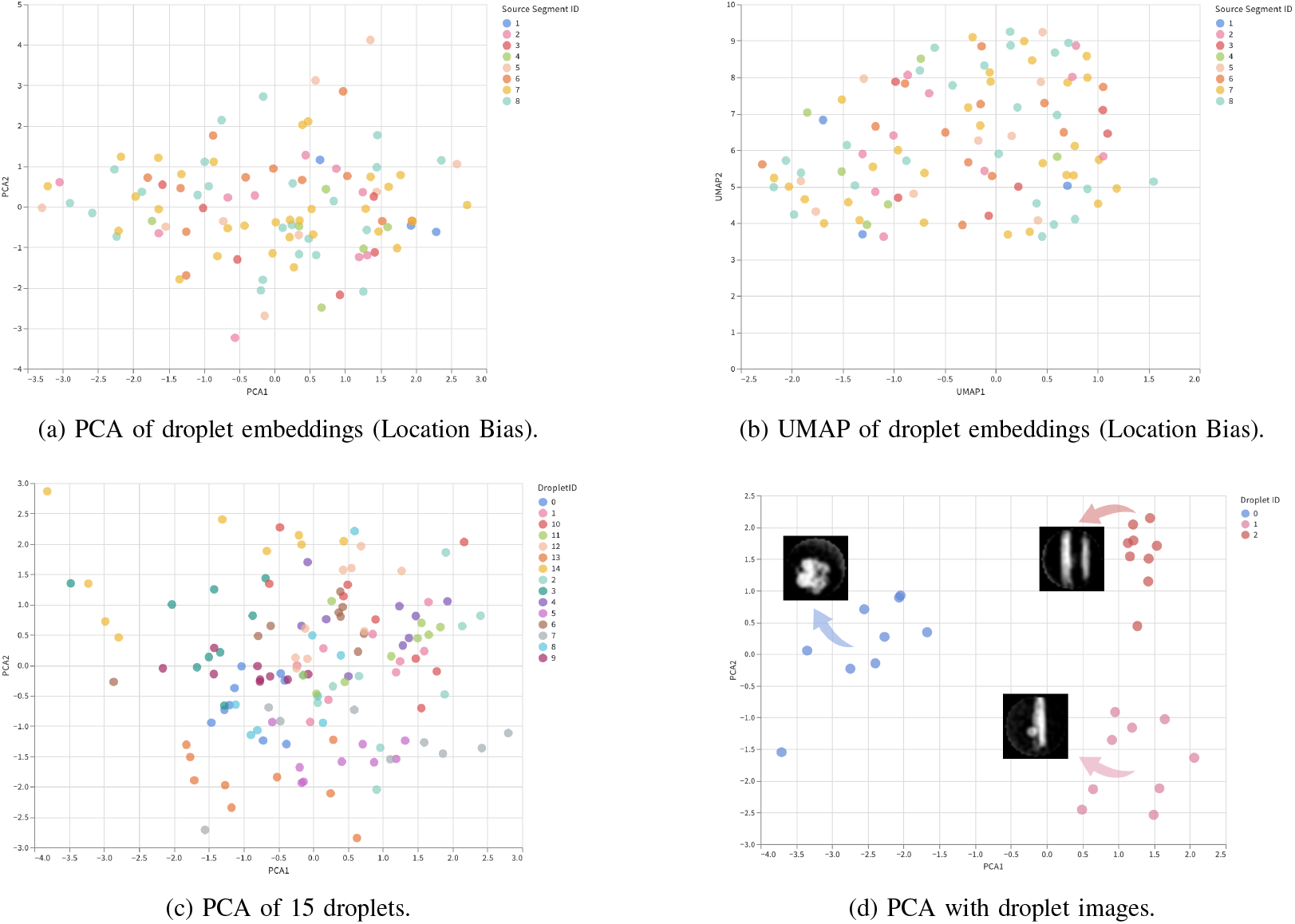
Dimension reduction of droplet embeddings of 8 droplets from different segments of the image using PCA **(a)** and UMAP **(b)** and for 15 droplets using PCA **(c)**. PCA of 3 droplet embeddings from the same droplets **(d)** with corresponding droplet image visualized. Data corresponds to Figure 4.

**Supplementary Figure 6:**
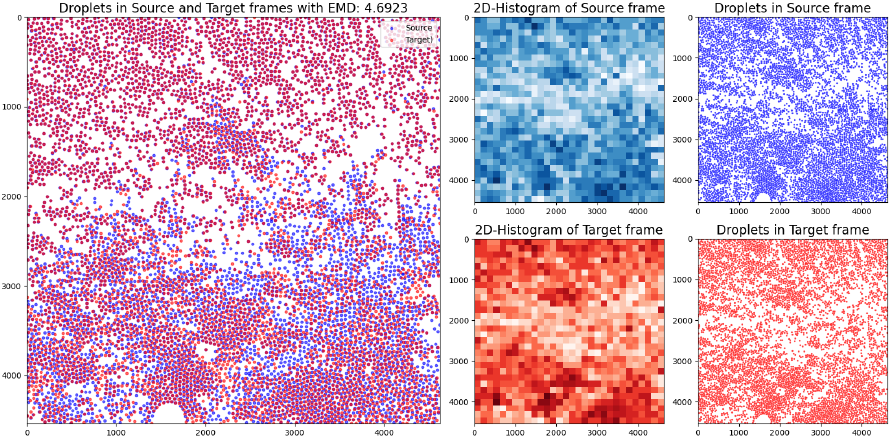
**Left:** Overview of the distribution of source (red) and target (blue) droplets from time-lapse series images for EMD calculation. **Middle:** 2D-Histogram of Droplet Locations at source (blue) and target (red). **Right:** Individual source and target Droplets separated.

We evaluated whether visual embeddings could pick up some image artifacts related to location. Such an analysis is important, as biases stemming from droplet position could influence final tracking outcome. All our data used to train and evaluate original embeddings are denoted as “low movement”. Hence, we could have potentially mistaken the exploitation of such local artifacts for good performance. To confirm this is not the case, we visualized the influence o f t he original droplet location on the embedding space. We split a single frame of the original image into segments and colored droplets based on the segment they are located in. In Supplemental Figure 5 we observed that the embeddings of droplets coming from different segments are well mixed for both reduction algorithms. Moreover, if we compare the UMAP and PCA visualizations from Supplemental Figure 5a,b and c with Supplemental Figure 5d, we could show that embeddings coming from data assigned to the same segment do not form similar structures to the ones coming from the same droplet across frames. This indicates that visual features rather than location and position are affecting dimensional reduction clustering.

## Appendix VII

### Distance Metric for image classification

To classify the movement between two frames, we propose a distance metric based on the Earth Movers Distance (EMD), which calculates the total amount of work (distance) needed to move all elements of a source distribution (frame) to a destination distribution^3^. It is calculated as follows:

1. The positions of the droplets are normalized by the median droplet radius.
2. Segment the source and the target frames into bins (in our case a gird with 30×30 bins) and count the number of droplets in each bin, then divide by the total number of droplets in the frame (both the source and the target frame individually).
3. Calculate the EMD between the histogram distributions.
4. For the classification for the full video take the median of all steps.

Values below 1 are classified as small movement; values between one and two are classified as medium movement, and values above two are classified as large movement. Implementation of EMD classification is located in the code base. We have obtained EMD metrics for all images analyzed by the algorithm and obtained median, mean and standard deviation of EMD metrics for each image. We calibrated the median EMD against computationally moved droplet images to produce *µ*m and pixel distance. To compare median EMD with actual distances, EMD *>* 0.4 corresponds to an average droplet movement of *>* 130 *µ*m or *>*2 droplet diameter per time-interval (Supplemental Figure 10).

**Supplementary Figure 7:**
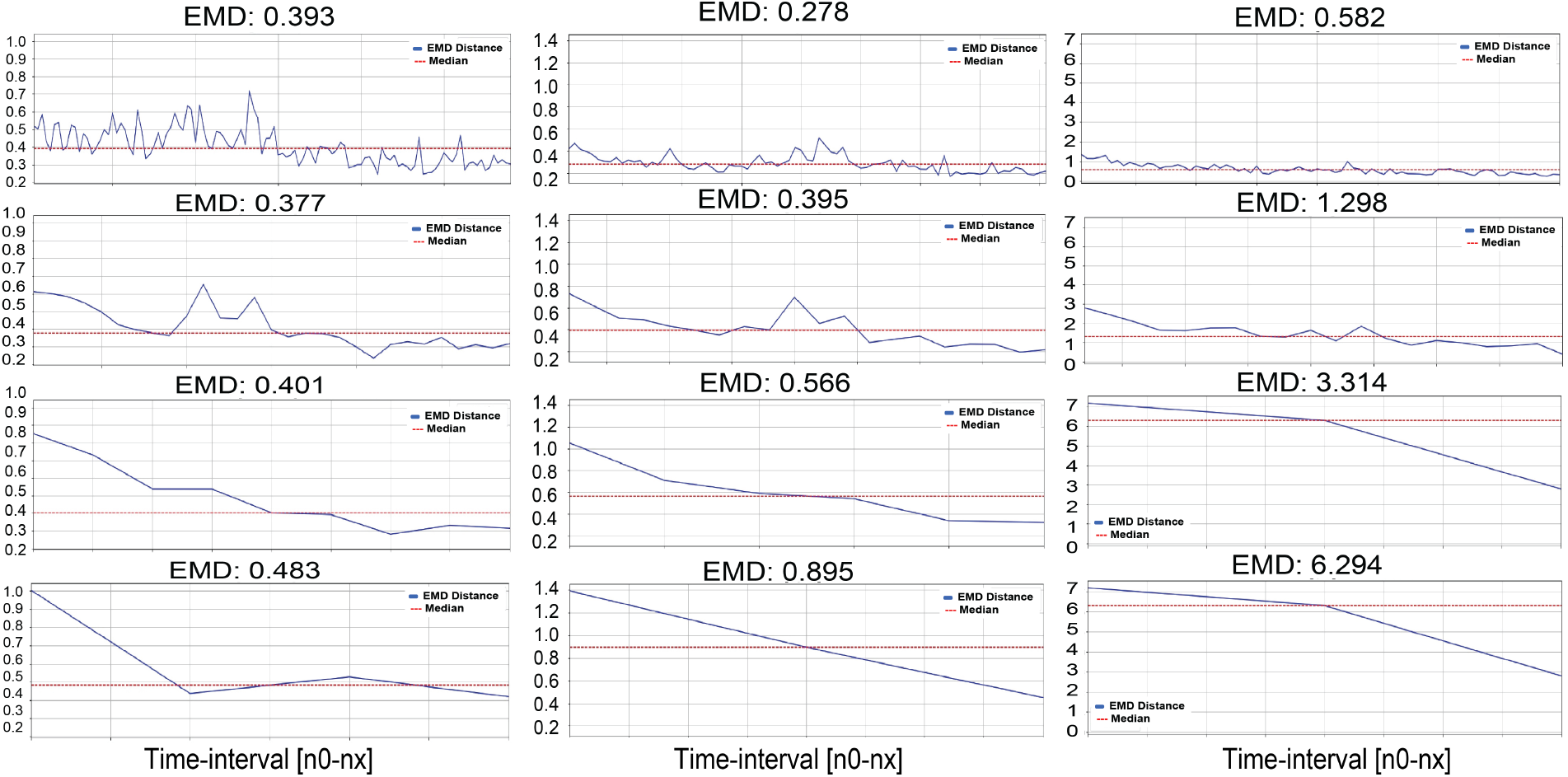
Raw data of EMD distance measurements for Dataset 1, corresponding to Table III (and 6). The EMD values are plotted as line plots at each time transition for low (left), medium (middle) and large (right) movement at 1 min, 5 min, 15 min, and 30 min time resolution (top to bottom). Median EMD displayed as red line and stated above.

In Appendix II-A and II-C, the raw experimental images are described in detail. Supplemental Figure 8 shows bar plots of the EMD distance median for Dataset 1 for low, low-to-medium, medium and large movement at the different, sampled time resolutions (corresponding to data in Table III. Supplemental Figure 9 shows the raw EMD values as lineplots for Dataset 3, time-lapse series without visual features (nanoparticles) for low-to-medium movement (top) and high movement (bottom) (used in Figure 7). Raw EMD values of Dataset 1 are plotted in Supplemental Figure 7 as line plots for low, medium and large movement at the different, sampled time resolutions (used in Table III and Figure 6).

**TABLE III:** Summary of metrics including EMD Distance, tracking accuracy, and droplet average movement. **EMD Distance**: Median EMD across whole time series. **Average movement:** Total image movement in droplet diameter and micrometer (µm) for 30 min intervals (converted from EMD, Supplemental Figure 10). **Percent trackable droplets:** calculated from DropMap analysis (in Matlab [5]). **Accuracy:** Metric from tracking evaluation and parameter analysis. Trajectory accuracy output from tracking at different temporal resolutions (5 min, 15 min and 30 min). **Percent Droplet [%]:** Tracking success defined as frequency of droplets with OT-scores of all transitions *x*_*t*_ to *x*_*t*+1_ *>* 0.5.

| Metric | Small Movement | Medium Movement | Large Movement |
| --- | --- | --- | --- |
| EMD Distance | 0.483 | 0.895 | 6.294 |
| Average movement: in droplet diameter (dd) in $\mu\text{m}$ | 2dd<br>($\sim 126\mu\text{m}$) | 12dd<br>($\sim 800\mu\text{m}$) | 160dd<br>($\sim 10,000\mu\text{m}$) |
| [%] previously possible to track using Matlab script [5] | 23.2% | 0.0% | 0.7% |
| Accuracy: 5 Min | 0.793 | 0.864 | 0.845 |
| Accuracy: 15 Min | 0.692 | 0.823 | 0.681 |
| Accuracy: 30 Min | 0.682 | 0.750 | 0.617 |
| Percent Droplets [%] with OT scores $> 0.5$ | 91.6% | 99.3% | 61.8% |

**Supplementary Figure 8:**
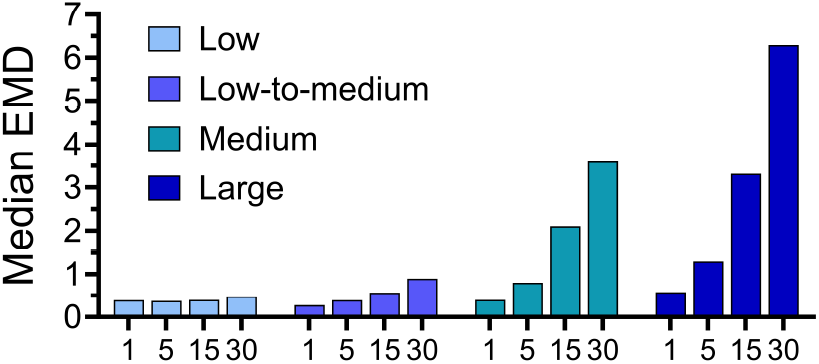
Histogram of median EMD distance for Dataset 1 at different, sampled time resolutions. Line plots for all bars are visualized in Supplemental Figure 7. (Low-to-medium movement was not included into tracking output of Table III, but showed similar tracking qualities).

**Supplementary Figure 9:**
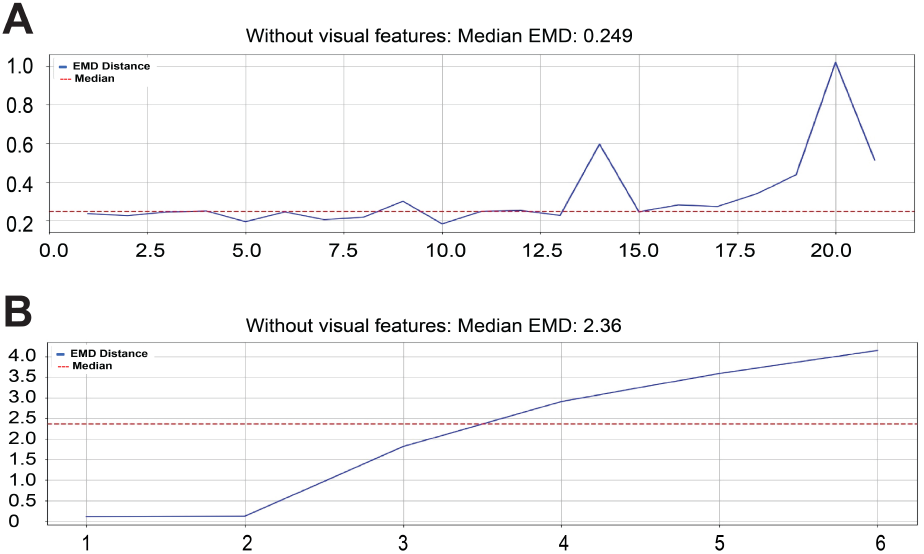
Raw data of EMD distance measurements of Dataset 3 used in Figure 7. The EMD values are plotted as line plots at each time transition. Median EMD displayed as red line and stated above. **A** low-to-medium movement **B** medium-to-large movement

**Supplementary Figure 10:**
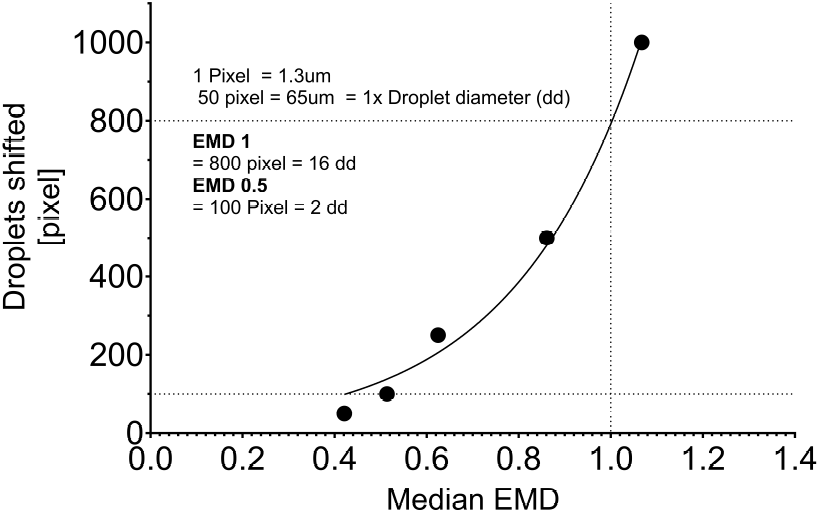
Calibration of median Earth Mover distance from images with defined, computationally introduced movement. Droplets were shifted by steps of 50 pixel and each shift saved with its original, source frame. Pixel shift was converted to *µ*m and droplet diameter. Median EMD was calculated for all images and plotted against the droplet shift. EMD to Pixel correlation was modeled with Exponential Growth Equation fit.

## Appendix VIII

### High-temporal resolution data barcode visualization

Droplet barcoding consists of encapsulation of droplets with a fluorescent d yes a t a s pecific co ncentration (e .g. Calcein-Green). Separate samples of droplets with blue, green and orange dyes were encapsulated, mixed and analzyed. Fluorescent dyes display stable intensities across a time-lapse experiment. Channel specific fi ltering (*>* 10 .000 fo r DA PI, ab ove 3000 for FITC and above 500 for TRITC) generates droplet IDs and data for color barcodes for each channel.

To indicate “failure” to track, mean droplet intensities were plotted. Minor changes in fluorescent i ntensity a re expected due to low bleaching effects or changes in contrast. However, for intensity changes larger than *>*20% relative to the average across all droplets was considered a false trajectory at that interval. Cross-validating high-temporal resolution data resulted in almost perfect trajectories (Supplemental Figure 11). However, some events remain incorrectly tracked (approximately 10-15 out of 4000 events), most likely due to fluorescent aggregates or residual (very-fast) movement. Nevertheless, most (*>* 95%) of droplets were detected with no or only marginal fluorescence changes confirming high-temporal resolution data.

**Supplementary Figure 11:**
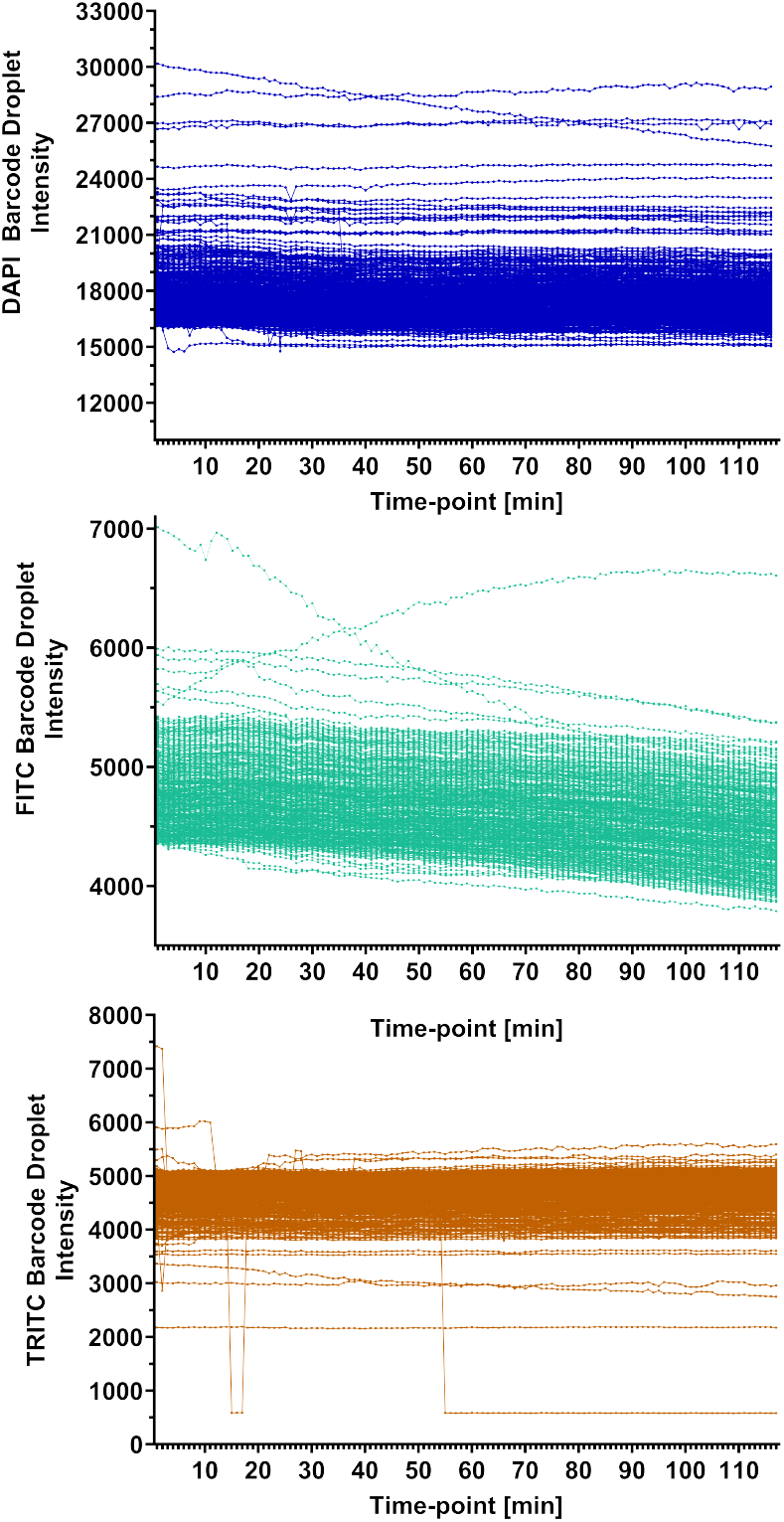
Raw data from high-temporal resolution fluorescence barcoding. Droplet barcodes were calculated from 1.25 min time-lapse series data. For each fluorescent barcode channel, 500 droplets with the highest per-channel intensities were filtered to avoid overlap (DAPI = blue, FITC = green, and TRITC = orange). Mean intensities were plotted against each time-point.

## Footnotes

1 The optimal transport problem is called unbalanced, if the source mass is not equal to the target mass. Unbalanced OT refers rebalancing or removing some mass from the problem by relaxing the marginal conditions. Since not all droplets can be transported to the target frame at all times (i.e., more droplets in the source than in the target frame) or vice versa not all droplets in the target have a droplet in the source frame, this requires computational adaption [27].

2 The entropy-regularizer is defined as: *E*(*T*) = Σ_*i,j*_ *T*_*i,j*_ log *T*_*i,j*_

3 Apart from the detection of droplets, this metric is fully independent of the algorithm or tracked distances.

